# Mutations from bat ACE2 orthologs markedly enhance ACE2-Fc neutralization of SARS-CoV-2

**DOI:** 10.1101/2020.06.29.178459

**Authors:** Huihui Mou, Brian D. Quinlan, Haiyong Peng, Yan Guo, Shoujiao Peng, Lizhou Zhang, Meredith E. Davis-Gardner, Matthew R. Gardner, Gogce Crynen, Zhi Xiang Voo, Charles C. Bailey, Michael D. Alpert, Christoph Rader, Hyeryun Choe, Michael Farzan

## Abstract

The severe acute respiratory syndrome coronavirus 2 (SARS-CoV-2) spike (S) protein mediates infection of cells expressing angiotensin-converting enzyme 2 (ACE2). ACE2 is also the viral receptor of SARS-CoV (SARS-CoV-1), a related coronavirus that emerged in 2002-2003. Horseshoe bats (genus *Rhinolophus*) are presumed to be the original reservoir of both viruses, and a SARS-like coronavirus, RaTG13, closely related SARS-CoV-2, has been isolated from one horseshoe-bat species. Here we characterize the ability of S-protein receptor-binding domains (RBDs) of SARS-CoV-1, SARS-CoV-2, and RaTG13 to bind a range of ACE2 orthologs. We observed that the SARS-CoV-2 RBD bound human, pangolin, and horseshoe bat (*R. macrotis)* ACE2 more efficiently than the SARS-CoV-1 or RaTG13 RBD. Only the RaTG13 RBD bound rodent ACE2 orthologs efficiently. Five mutations drawn from ACE2 orthologs of nine *Rhinolophus* species enhanced human ACE2 binding to the SARS-CoV-2 RBD and neutralization of SARS-CoV-2 by an immunoadhesin form of human ACE2 (ACE2-Fc). Two of these mutations impaired neutralization of SARS-CoV-1. An ACE2-Fc variant bearing all five mutations neutralized SARS-CoV-2 five-fold more efficiently than human ACE2-Fc. These data narrow the potential SARS-CoV-2 reservoir, suggest that SARS-CoV-1 and -2 originate from distinct bat species, and identify a more potently neutralizing form of ACE2-Fc.

## INTRODUCTION

Coronaviruses are enveloped positive-strand RNA viruses of the family *Coronaviridae* (Cui et al., 2019). At least seven coronaviruses infect humans. Four of these, HCoV-229E, -OC43, -HKU1, and -NL63 cause mild upper respiratory tract symptoms in most infected persons. In contrast MERS-CoV and SARS-CoV-1 cause severe, often fatal, infections. The recently emerged SARS-CoV-2 is closely related to SARS-CoV-1, and causes the sometimes fatal disease COVID-19. SARS-CoV-2, like SARS-CoV-1, uses the receptor ACE2 to infect cells (Li et al., 2003). Association with ACE2 and infection of ACE2-expressing cells is mediated by the SARS-CoV-1 and SARS-CoV-2 spike (S) proteins. These S proteins are type I viral entry proteins similar to influenza hemagglutinin and the HIV-1 envelope glycoprotein (Li, 2016). Like these latter entry proteins, the S protein is processed into two domains, S1 and S2 (Walls et al., 2020; Wrapp et al., 2020), either in the virus-producing cell (SARS-CoV-2) or in the ACE2-expressing target cell (SARS-CoV-1). S1 binds ACE2, whereas S2 anchors the S protein to the viral membrane and mediates fusion with the target-cell membrane. Somewhat unusually for type I entry proteins, the S1 domains of SARS-CoV-1 and -2 include distinct, independently folded receptor-binding domains (RBDs) of approximately 200 amino-acids (Li et al., 2005a; Wong et al., 2004). The RBD is the primary neutralizing epitope on the SARS-CoV-2 S protein.

In addition to these human viruses, a number of SARS-like viruses have been isolated from horseshoe bats (genus *Rhinolophus*) (Graham and Baric, 2010; Li et al., 2005b; Menachery et al., 2015). These bats are also the presumed long-term reservoirs of SARS-CoV-1 and -2, but it remains unclear exactly which *Rhinolophus* species serves as the most recent bat host of each virus. Although some SARS-like viruses have deletions in their receptor-binding domains (RBDs), likely precluding their use of ACE2, other SARS-like viruses, like SARS-CoV-1 and -2, have intact RBDs and retain their ability to bind and enter cells through ACE2 (Li et al., 2005a; Li et al., 2006a; Li et al., 2005b; Lu et al., 2020). One such SARS-like virus, RaTG13, isolated from the horseshoe bat species *R. affinis* is closely related to SARS-CoV-2 (96.2% nucleotide identity) (Zhou et al., 2020).

Abundant data implicate the palm civet as a reservoir intermediate of SARS-CoV-1 (Li et al., 2006a). For example, the palm-civet ACE2 ortholog is an efficient receptor for SARS-CoV-1 (Li et al., 2005c). Also, SARS-CoV-1 has been isolated from palm civets at an exotic animal market in the Guangdong province, where the virus first infected humans in 2002 (Guan et al., 2003). Finally, a second independent human transmission of SARS-CoV-1, which occurred in the winter of 2003, was directly traced to a restaurant serving palm civets (Wang et al., 2005). Analogously, the pangolin has been suggested to be a reservoir intermediate for SARS-CoV-2, and a closely related SARS-like virus that utilizes ACE2 has been isolated from this species (Xiao et al., 2020). However these pangolins showed signs of coronaviral disease, more consistent with an intermediate host than a long-term reservoir. The near identity of the RBD from this pangolin coronavirus and the SARS-CoV-2 RBD has suggested that SARS-CoV-2 arose from a recombination event between a pangolin- and a bat-derived coronavirus, although it remains undetermined in what species this putative recombination event occurred (Xiao et al., 2020; Zhang et al., 2020b).

Soluble forms of ACE2, including its immunoadhesin form ACE2-Fc (also described as ACE2-Ig), neutralize both SARS-CoV-1 and SARS-CoV-2 (Hoffmann et al., 2020; Moore et al., 2004; Walls et al., 2020). ACE2-Fc may be useful for treating infected persons, and, in the absence of an effective vaccine, it might protect individuals from an initial infection (Monteil et al., 2020). Here we characterize nine orthologs of ACE2, including those of pangolin and two horseshoe bat species, for their ability to bind the SARS-CoV-1, SARS-CoV-2, and RaTG13 RBD. We observed that the SARS2-CoV-2 RBD (SARS2-RBD) binds human, pangolin and horseshoe-bat (*R. macrotis*) ACE2 more efficiently than the SARS-CoV-1 RBD (SARS1-RBD). In contrast the SARS1-RBD bound mouse, palm civet, pig, and dog ACE2 orthologs more efficiently. A subset of residues drawn from nine horseshoe-bat ACE2 orthologs enhanced human ACE2 binding to the SARS2-RBD and neutralization of SARS-CoV-2 S-protein pseudotyped retroviruses (SARS2-PV). These data indicate that SARS-CoV-2 S-protein has not fully adapted to human ACE2 and thus elements of its reservoir species ACE2 ortholog can be used to improve the neutralization potency of ACE2-Fc.

## RESULTS

### SARS2-RBD binds more efficiently to human, pangolin and *R. pusillus* ACE2 orthologs than does SARS1-RBD

ACE2 utilization has been predictive of the susceptibility of a species to SARS-CoV-1 infection and provided useful insight to the zoonotic origins of this virus (Li et al., 2006b; Li et al., 2005c). We therefore initiated similar studies of SARS-CoV-2. We first investigated the ability of immunoadhesin forms of the SARS1-RBD, SARS2-RBD, MERS-CoV RBD (MERS-RBD) and RaTG13-RBD, each fused to the Fc domain of human IgG1 (**Figure S1A**) to bind HEK293T cells expressing the ACE2 orthologs of nine species. These species include palm civet, a known reservoir intermediate of SARS-CoV-1, and pangolin, a proposed reservoir intermediate of SARS-CoV-2 (Guan et al., 2003; Li et al., 2005c; Xiao et al., 2020). We also characterized the ACE2 orthologs of two horseshoe bat species (*R. macrotis, R. pusillus*), selected because they are closely related to *R. affinis*, the source of RaTG13, but in contrast to *R. affinis*, their ACE2 orthologs have been described. Finally we characterized human, pig, dog, mouse, and rat ACE2 orthologs to evaluate their potential as reservoirs or model systems for SARS-CoV-2.

Each ACE2 ortholog was transiently expressed on HEK293T cells. Expression of each ortholog was quantified by flow cytometry using an antibody recognizing myc tag fused to the amino terminus of each ortholog (**Figure S1B**). Flow cytometry was also used to measure the relative ability of each RBD-Fc variant to bind these ACE2 orthologs. Histograms in **Figure 1** show the results of these studies and their associated bars quantify RBD binding (mean-fluorescence normalized to ACE2 expression). We observed that SARS2-RBD-Fc bound cells expressing human, pangolin, *R. macrotis* ACE2 more efficiently than did SARS1-, RaTG13-, or MERS-RBD-Fc. The SARS2-RBD-Fc also efficiently bound pig and dog ACE2, although less efficiently than the SARS1-RBD-Fc. The SARS1-RBD-Fc efficiently bound palm civet ACE2, as we have previous reported. Notably the RaTG13-RBD-Fc bound both rat and mouse ACE2 orthologs more efficiently than any other RBD-Fc. The inability of the SARS2-RBD to bind murine ACE2 suggests that, in contrast to SARS-CoV-1, wild-type mice are unlikely to support useful replication of SARS-CoV-2 (Li et al., 2004; Roberts et al., 2007; Subbarao et al., 2004). Also of note, no RBD-Fc bound the *R. pusillus* ACE2 ortholog, indicating that this horseshoe bat is not a reservoir for either SARS-CoV virus. As expected, the MERS-RBD-Fc control construct only bound cells expressing DPP4 (CD26) the MERS-CoV receptor (Raj et al., 2013). These data are consistent with the hypothesis that pangolin is a reservoir intermediate for SARS-CoV-2 and suggest that the bat reservoir for this virus has an ACE2 with an S-protein binding site more similar to *R. macrotis* ACE2 than to *R. pusillus* ACE2.

**Figure 1.**
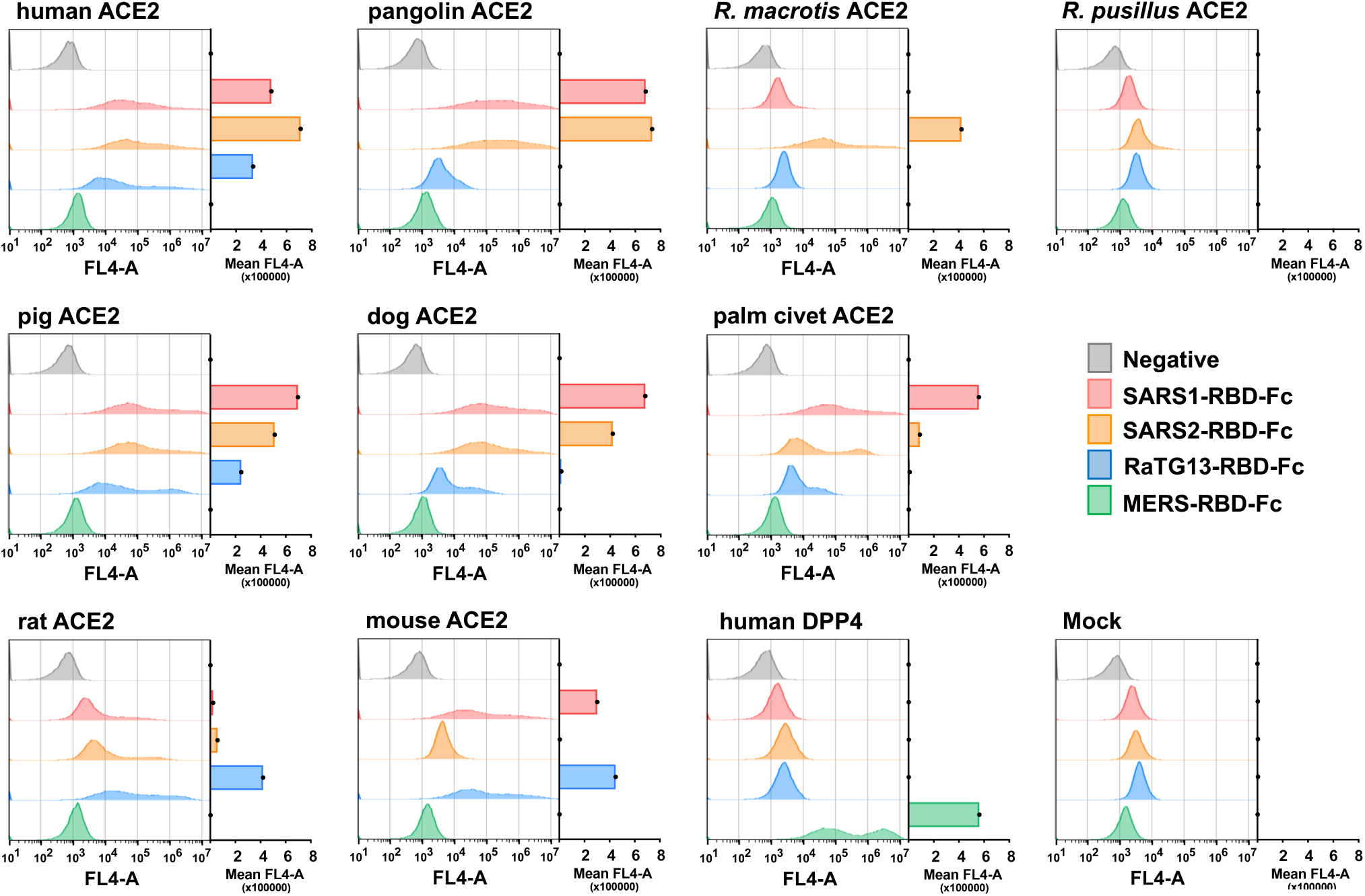
The receptor-binding domains (RBD) of SARS-CoV-1, SARS-CoV-2, and RaTG13 S proteins bind differentially to ACE2 orthologs. HEK239T cells transfected to express the ACE2 orthologs of the indicate species, human DPP4 (the receptor of MERS-CoV), or with vector alone (mock) were assayed by flow cytometry for their ability to bind SARS1-RBD-Fc, SARS2-RBD-Fc, RaTG13-RBD-Fc, or MERS-RBD-Fc. Histograms display a representative experiment, and bars display the results of two experiments normalized to ACE2 expressing levels shown in Figure S1B and determined by an antibody recognizing an amino-terminal myc-tag.

### Quantitative binding studies of SARS1- and SARS2-RBD-Fc to human, pangolin and *R. macrotis* ACE2 orthologs

To better quantify the association between these RBDs and human, pangolin, and *R. macrotis* ACE2 orthologs, we used surface plasmon resonance (SPR) to measure association- or on-rates (*k*_*on*_), dissociation or off-rates (*k*_*off*_), and equilibrium dissociation constants (*K*_*d*_) of these ligands. Soluble, monomeric forms of human ACE2, an enzymatically inactive human-ACE2 variant (ACE2-NN), pangolin ACE2, and *R. macrotis* ACE2 were purified from supernatants of transfected Expi239 cells (**Figure S1A**) and analyzed. Each RBD-Fc variant was captured with a mouse anti-human IgG CH2 monoclonal antibody immobilized on a CM5 sensor chip and monitored for binding to serially diluted soluble-monomeric ACE2 variants, as represented in **Figure 2A**. We observed that the SARS2-RBD-Fc bound monomeric ACE2 two- to three-fold more efficiently than the SARS1-RBD, consistent with previous reports (Wrapp et al., 2020), with a *K*_*d*_ of approximately 100 nM (**Figure 2B**). Both RBD-Fc bound pangolin ACE2 less efficiently, with SARS1- and SARS2-RBD-Fc binding with approximate *K*_*d*_ of 1.5 and 1 µM, respectively. Only SARS2-RBD-Fc bound detectably to *R. macrotis* ACE2, with a *K*_*d*_ of 4.6 µM. As expected, RaTG13-RBD-Fc and MERS-RBD-Fc bound with affinities greater than 35 µM or undetectably. A comparison of binding kinetics (**Figure 2C**) shows that SARS1-RBD has a faster on-rate but also faster off-rate for human ACE2 and human ACE2-NN than does the SARS2-RBD, suggesting that the latter RBD may be less ordered in solution. Notably SARS2-RBD-Fc associates more quickly with pangolin ACE2 than with human ACE2, but its off-rate from the pangolin ortholog is markedly faster, resulting a net lower affinity for this ACE2 ortholog. Thus small differences in the ability of human and pangolin ACE2 orthologs to support SARS2-PV infection, as shown in Figure 1, can mask relatively large affinity differences, presumably because the S protein engages multiple ACE2 receptors on the cell surface.

**Figure 2.**
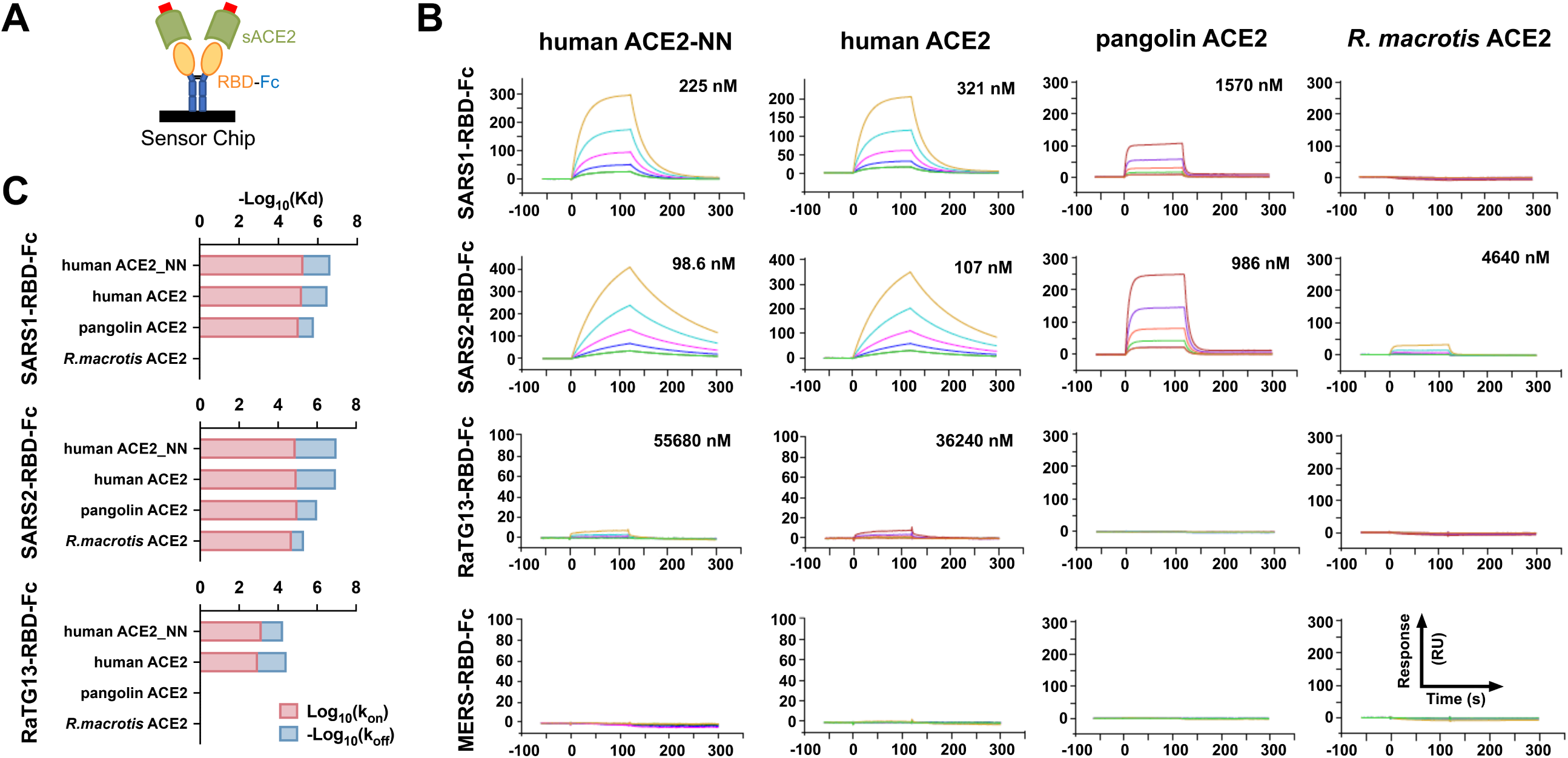
Quantitative comparisons of binding of RBD-Fc variants to soluble monomeric ACE2 orthologs. (**A**) Biacore surface plasmon resonsance (SPR) studies were performed the indicated RBD-Fc (yellow and blue) captured on the sensor chip. The soluble monomeric forms of the indicated ACE2 variants (green) were injected for analysis as represented in the figure. Red indicates a four-amino acid at tag at the ACE2 C-termini. (**B**) Biacore X100 sensorgrams representing results from studies in which the indicated RBD-Fc variants were captured by an anti-human Fcγ antibody immobilized on a CM5 chip after instantaneous background depletion. Monomeric forms of human ACE2, human ACE2 lacking its active-site histidines (ACE2-NN), pangolin ACE2, or bat (*R. macrotis*) ACE2 were injected at five concentrations, serially diluted by two-fold from the highest concentration (100 nM for human ACE2 variants, 500 nM for non-human orthologs). The experiment is representative of two with similar results. (**C**) The association rate constant (*k*_*on*_), dissociation rate constant (*k*_*off*_), and equilibrium dissociation constant (*K*_*d*_) were calculated and plotted from the study shown in (A). The negative of log_10_ (*K*_*d*_) is represented as the sum of log_10_ (*k*_*on*_), shown in red, and the negative of log_10_ (*k*_*off*_), shown in blue. Figure at bottom indicates that the RBD-Fc is immobilized and the monomeric soluble ACE2 is injected in this study.

### SARS2-RBD-Fc and ACE2-Fc efficiently neutralize SARS1- and SARS2-S-protein mediated entry

The RBD-Fc and ACE2 variants studied in Figure 2 were also compared in **Figure 3** for their ability to neutralize retroviral particles pseudotyped with the SARS-CoV-1 S protein (SARS1-PV), the SARS-CoV-2 S protein (SARS2-PV), or the G protein of vesicular stomatitis virus (VSV-G-PV). To test inhibition by these RBD-Fc, cells were preincubated with the indicated protein before pseudovirus addition. SARS1-RBD-Fc and SARS2-RBD-Fc efficiently inhibited SARS1-PV and SARS2-PV, but not VSV-G-PV (**Figure 3A**). The SARS2-RBD-Fc inhibited both SARS pseudoviruses more efficiently than SARS1-RBD-Fc. Expectedly neither the MERS-RBD-Fc nor the RaTG13-RBD inhibited any pseudovirus. In addition, soluble monomeric ACE2 (sACE2) proteins were tested for their ability to neutralize pseudoviruses. Consistent with their affinities, monomeric human ACE2 and ACE2-NN neutralized SARS2-PV more efficiently than SARS1-PV, whereas no neutralization was observed with soluble pangolin or *R. macrotis* ACE2 (**Figure 3B**). Soluble human ACE2 fused to a human IgG1 Fc domain (ACE2-Fc) neutralized SARS1-and SARS2-PV approximately 10-fold more efficiently than monomeric ACE2, suggesting that both arms of this dimeric ACE2-Fc construct simultaneously engaged distinct RBD regions on the pseudovirion. These latter observations suggest that ongoing efforts to use monomeric ACE2 to treat COVID-19 infection (Zhang et al., 2020a) will be less effective than treatments using ACE2-Fc.

**Figure 3.**
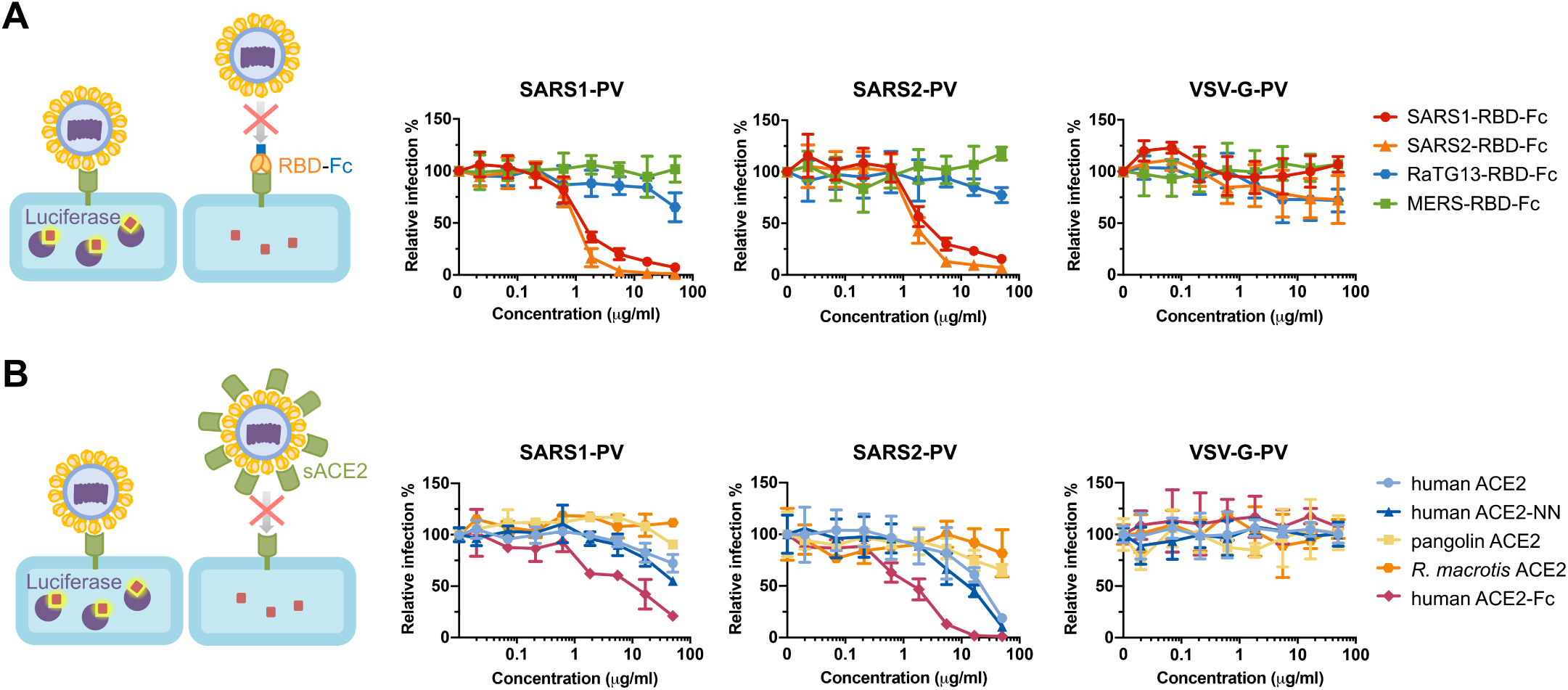
SARS1-RBD-Fc, SARS2-RBD-Fc, and human ACE2-Fc inhibit S-protein-mediated infection. (**A**) The indicated concentrations of the indicated RBD-Fc variants were incubated with retroviral pseudoviruses (PV) pseudotyped with the S proteins of SARS-CoV-1 (SARS1-PV), SARS-CoV-2 (SAR2-PV), or with the G protein of vesicular stomatitis virus (VSV-G-PV). Infection is normalized to that in the absence of inhibitors. (**B**) Experiments similar to those in (A) except that the indicated soluble monomeric ACE2 variants, or human ACE2-Fc were used to inhibit the indicated pseudoviruses. Error bars in (A) and (B) indicate standard error of the mean (S.E.M), and are representative at least two experiments with similar results.

### Residues derived from horseshoe-bat ACE2 orthologs enhance neutralization of SARS2-CoV pseudoviruses

The SARS1- and SARS2-RBDs bind a common region of ACE2 (indicated in yellow in **Figure 4A**) that varies among human ACE2, pangolin ACE2, and horseshoe-bat ACE2 orthologs (**Figure 4B**). We speculated that SARS-CoV-2 had not fully adapted to the use of the human receptor, and elements of bat and perhaps pangolin ACE2 might enhance the affinity of human ACE2 for the SARS2-RBD. To test this hypothesis, we altered eight residues of human ACE2-Fc to their analogues in one or more horseshoe-bat orthologs (red and green in **Figure 4A**). In addition to eight mutations based on this rationale (Q24E, T27K, H34S, N49D, N49E, M82N, N90D, Q325E), we further probed position 34 with residues present in other species (H34E, H34D, H34Y), and tested an interesting residue-354 difference between pangolin and human ACE2 (G354H). G354 is adjacent to K353, a residue critical to RBD association with ACE2.

**Figure 4.**
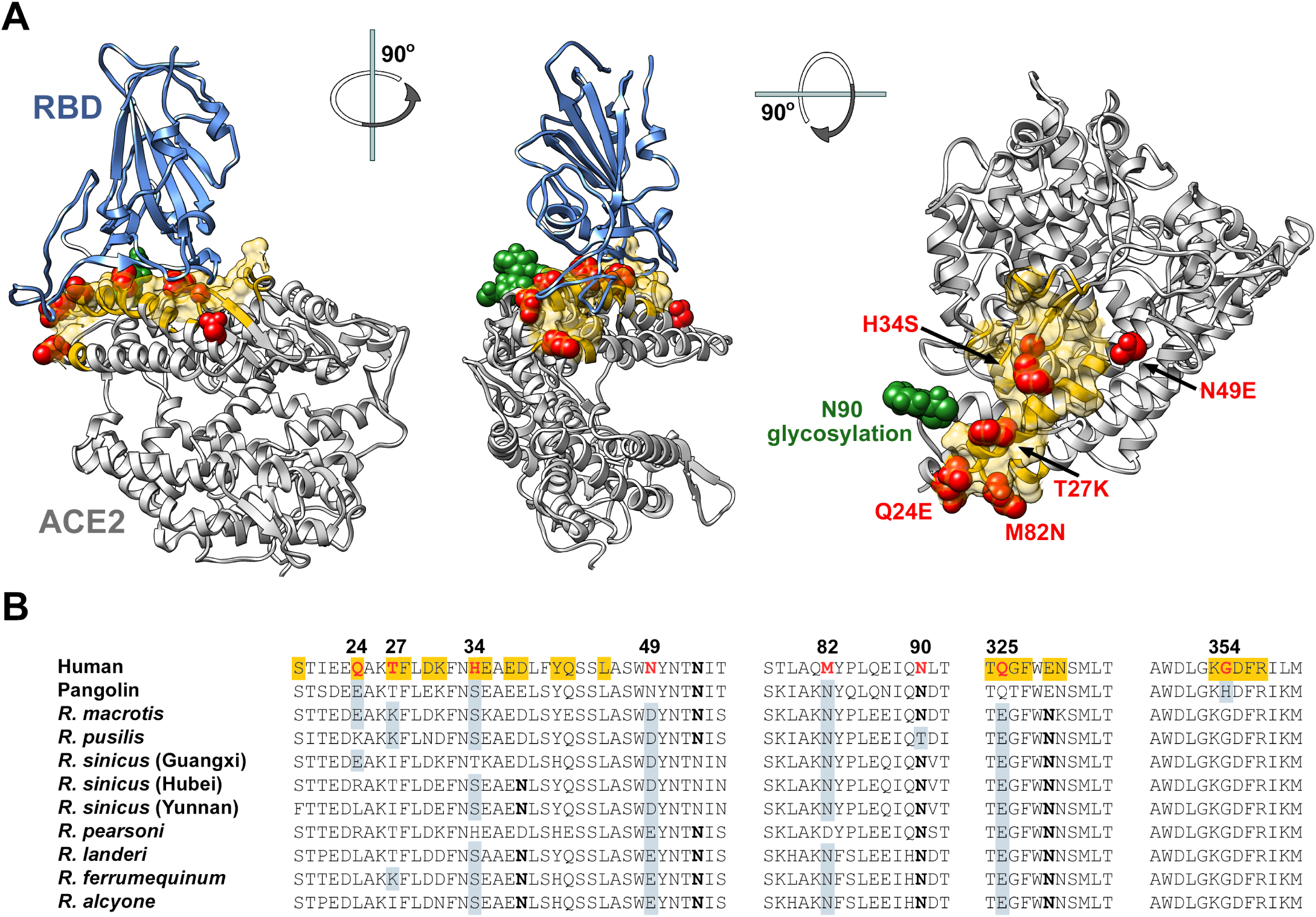
Horseshoe bat and pangolin-derived mutations characterized in Figures 5 and 6. **(A)** Human ACE2 (grey) is shown bound to the SARS-CoV-2 RBD (blue), based on PDB accession number 6M0J (Lan et al., 2020). Yellow indicates the ACE2 surface that is within 5.5 Å of the RBD. Red indicates residues that vary between human and horseshoe bat species and were characterized in the subsequent Figures. Specific mutations are indicated in the rightmost figure. A glycosylation at asparagine 90, also characterized, is indicated in green. Figure shows successive 90 degree rotations around the vertical and horizontal axes, respectively. The RBD is removed from the rightmost figure for clarity. (**B**) Sequences of human ACE2 directly associating, or proximal to, the SARS-CoV-2 RBD are shown, aligned with the sequence of the pangolin ACE2 ortholog, and those of all available horseshoe-bat ACE2 orthologs. Yellow indicates a residue that directly contacts the RBD. Red indicates human residues that were altered to the indicated pangolin and bat residues shown in gray and characterized in the subsequent figures. Bold indicates a glycosylated asparagine.

Human ACE2-Fc (hACE2-Fc) and these twelve variants were produced in Expi293T cells (**Figure S3**) and evaluated for their ability to neutralize SARS1-PV, SARS2-PV, and VSV-G-PV (**Figure 5A-C**). Seven of these variants neutralized SARS-CoV-2 more efficiently than wild-type hACE2-Fc: Q24E, T27K, H34S, H34D, N49E, M82N, and N90D. The ability of these ACE2-Fc variants to neutralized SARS2-PV was precisely reflected in their affinities for monomeric SARS2-RBD, as determined by SPR (**Figure 5D-E**). All variants that neutralized SARS2-PV more efficiently bound SARS2-RBD with higher affinity than wild-type ACE2-Fc, and all variants that neutralized less efficiently bound with lower affinity. These patterns were consistent regardless of whether ACE2-Fc (**Figure 5D; Figure S4**) or SARS2-RBD-Fc (**Figure S5**) was immobilized on the chip. Notably, G354H impaired SARS2-RBD association and SARS2-PV neutralization, and N90D, which removes a glycosylation at residue 90, markedly increased the SARS2-RBD on-rate.

**Figure 5.**
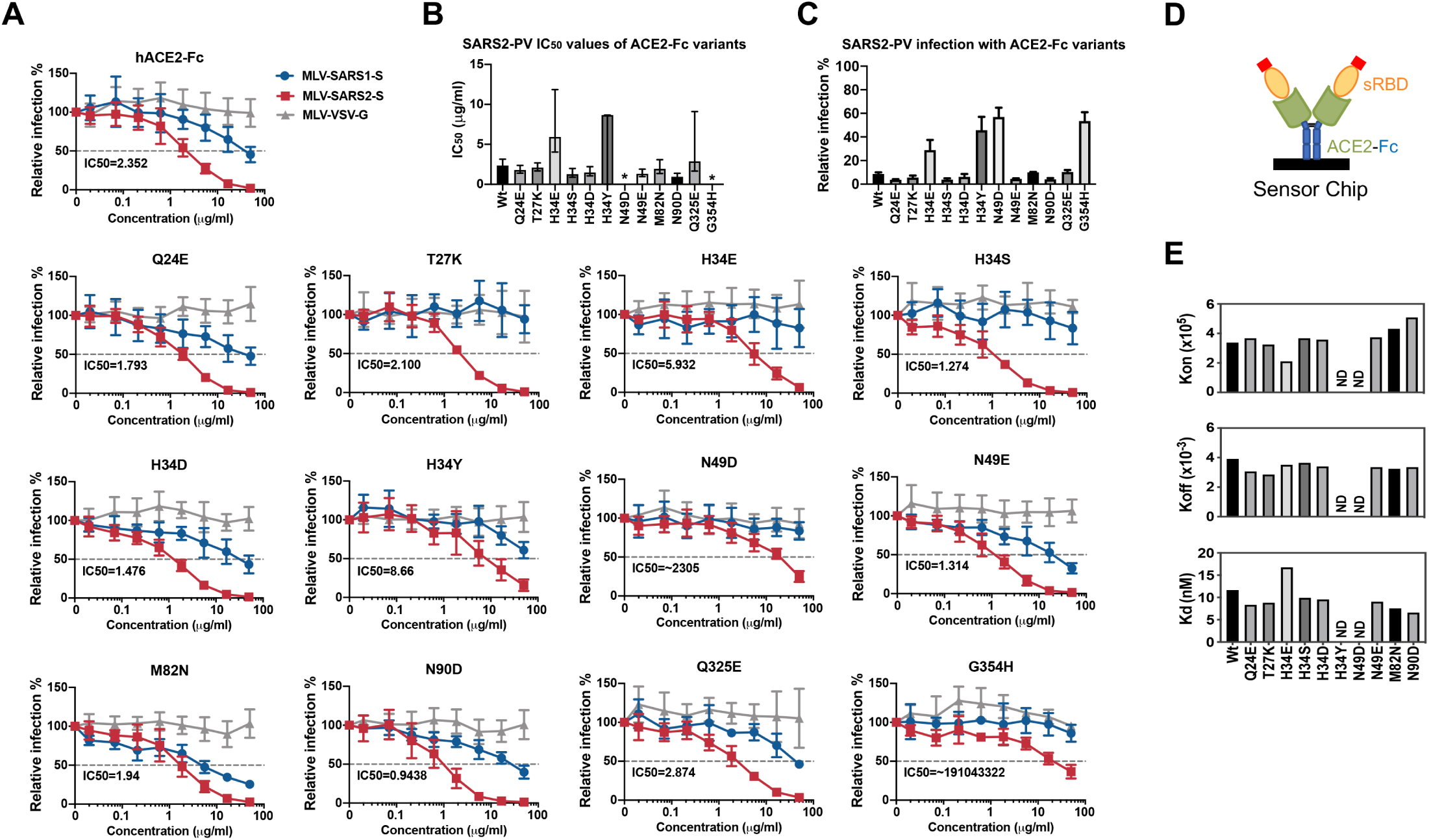
Residues from horseshoe-bat ACE2 orthologs improved human ACE2-Fc binding to SARS2-RBD and neutralization of SARS2-PV. (**A**) SARS1-PV (blue), SARS2-PV (red), or VSV-G-PV (grey) were used to infect ACE2-239T cells in the presence of the indicated concentration of human ACE2-Fc (hACE2-Fc) or hACE2-Fc with the indicated mutations. Mean half-maximal inhibitory concentration (IC_50_) shown each variant. Error bars indicate standard error of the mean (S.E.M) for replicates. Experiment is representative of at least two with similar results. (**B**) SARS2-PV IC_50_ values for three studies similar to those shown in (A) are plotted. Error bars indicate the upper and lower IC_50_ value observed for each hACE2-Fc variant. Asterisks indicate that the neutralization was incomplete and therefore no IC_50_ was calculated. (**C**) Percentage of SARS2-PV infection in the presence of 16.67 μg/ml indicated hACE2-Fc variant. Error bars indicate standard error of the mean (S.E.M). (**D**) A diagram is shown illustrating that hACE2-Fc variants were captured and that soluble monomeric SARS2-RBD was injected for these studies. (**E**) SPR analyses showing the *k*_*on*_, *k*_*off*_, and *K*_*d*_ of SARS2-RBD to hACE2-Fc variants.

We then selected five best-performing bat-derived changes for additional characterization alone (Q24E, T27K, H34S, N49E, and N90D) or combined (5M). These included two changes (Q24E; H34S) also present in pangolin ACE2. With the exception of T27K, each variant neutralized SARS2-PV more efficiently than wild-type hACE2-Fc (**Figure 6A-B; Figure S6**), with 5M neutralizing five-fold more efficiently (0.54 µg/ml, compared with 2.6 µg/ml for wild-type hACE2-Fc). Consistent with these neutralization studies, the 5M variant bound the SARS2-RBD with more than two-fold higher affinity (**Figure 6C-D**; 11.64 nM for wild-type hACE2-Fc; 5.454 nM for 5M). Thus individual mutations derived from horseshoe bats that enhance the neutralization potency of human ACE2-Fc can be combined to make an even more potent ACE2-Fc variant.

**Figure 6.**
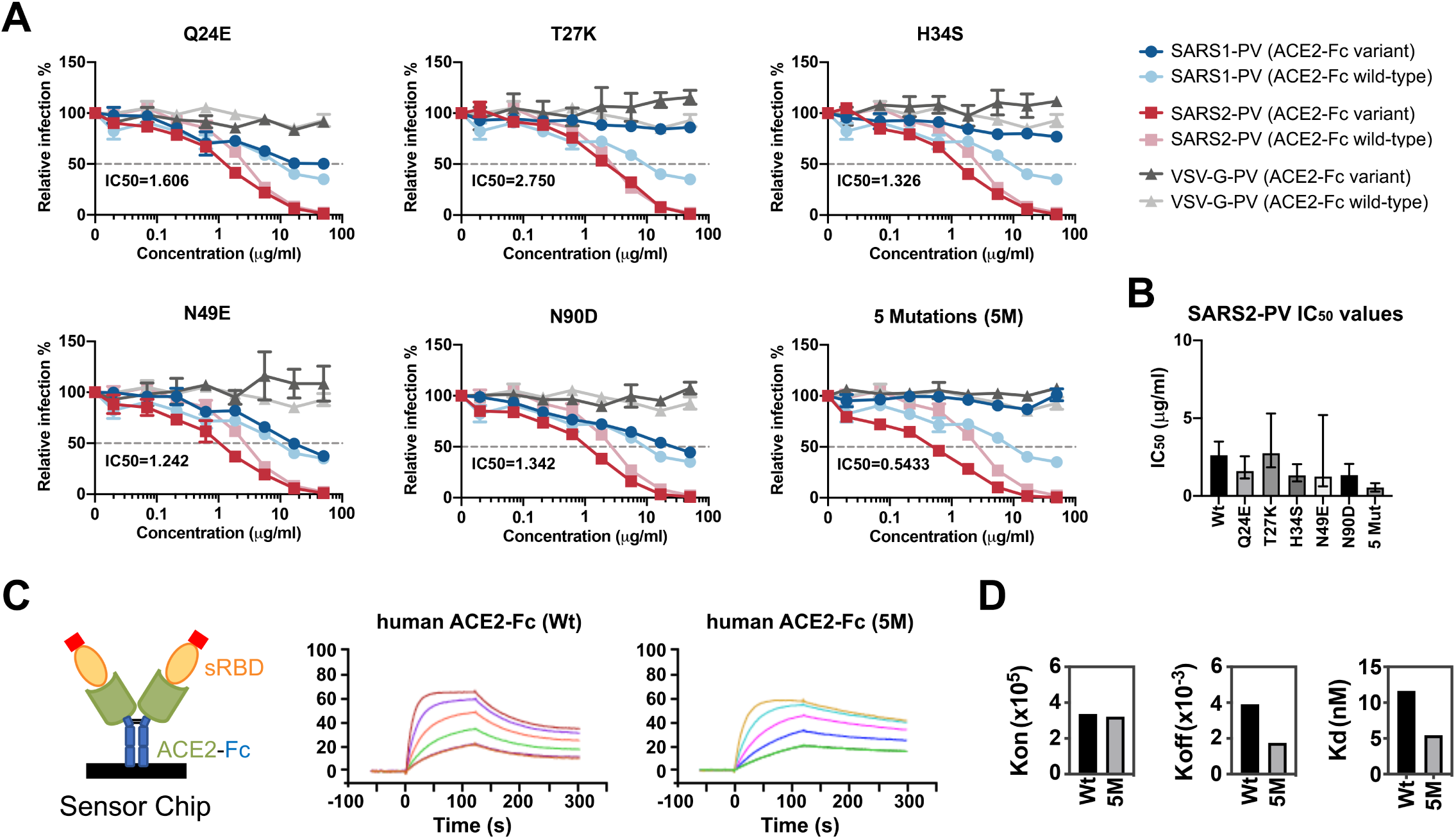
A combination of bat-derived residues improves SARS-PV neutralization five-fold. (**A**) Experiments similar to those shown Figure 5A except that the indicated ACE2-Fc variant (dark colors) is directly compared with wild-type human ACE2-Fc (light colors) for their ability to neutralize SARS1-PV (light and dark blue), SARS2-PV (light and dark red), or VSV-G-PV (light and dark grey). IC_50_ values for SARS-CoV-2 of the indicated ACE2-Fc variant over two independent studies are provided in bottom left of each figure. Neutralization results were normalized to the virus infection without any ACE2-Fc. Error bars indicate standard error of the mean (S.E.M). Neutralization data are representative of at least two experiments. (**B**) SARS2-PV IC_50_ values for two studies similar to those shown in (A) are plotted. Error bars indicate the upper and lower IC_50_ value observed for each hACE2-Fc variant. Asterisks indicate that the neutralization was incomplete and therefore no IC_50_ was calculated. (**C**) Sensorgrams of SPR analyses used to determine bind constants of the SARS2-RBD for wild-type hACE2-Fc (Wt) and hACE2-Fc with the five indicated mutations (5M). Diagram illlustrates that ACE2-Fc was captured in these studies while soluble monomeric RBD was injected. (**D**) *k*_*on*_, *k*_*off*_, and *K*_*d*_ of SARS2-RBD binding to wild-type hACE2-Fc or to hACE2-Fc 5M.

## DISCUSSION

Here we show that three ACE2 orthologs – human, pangolin, and *R. macrotis* bat – associate with the SARS2-RBD more efficiently than with the SARS1-RBD. In contrast, pig, dog, palm civet, and mouse orthologs bound more efficiently to SARS1-RBD, and neither RBD bound *R. pusillus* bat or rat ACE2. Unexpectedly the RBD of the SARS-like coronavirus RaTG13 isolated from horseshoe bats (*R. affinis*) did not bind either horseshoe bat ACE2 ortholog tested, indicating that the not-yet-described *R. affinis* ACE2 varies in significant ways from these horseshoe-bat orthologs. However, unlike the SARS2-RBD, the RaTG13-RBD bound both rat and mouse ACE2 orthologs efficiently. This observation might be used to generate viruses better adapted to these rodents, and perhaps providing an additional model of SARS-CoV-2 infection.

SARS2-RBD bound with lower affinity to *R. macrotis* ACE2 than to human ACE2, and it did not bind detectably to *R. pusillus* ACE2. This result suggests that SARS-CoV-2 emerged most recently from a different horseshoe-bat species. We hypothesized that the ACE2 ortholog of this hypothetical reservoir bat would be similar to the nine reported horseshoe bat ACE2 orthologs, and that the SARS-CoV-2 RBD would more efficiently bind ACE2 variants containing elements of these bat orthologs. We identified four such elements that enhanced ACE2-Fc neutralization of SARS2-PV and binding to the SARS2-RBD across multiple studies. Of interest, most of these changes impaired neutralization of SARS1-PV, implying that SARS-CoV-1 and SARS-CoV-2 derive from distinct horseshoe bat reservoirs. Two of these changes – Q24E and H34S – are notable because they are also present in pangolin ACE2 and are consistent with the proposal that the pangolin is a reservoir intermediate that facilitated SARS-CoV-2 adaptation to human ACE2. However, we show that G354H, a change unique to pangolin ACE2 among all sequenced ACE2 orthologs, impairs SARS2-RBD binding when it is introduced into human ACE2. It is worth noting that the pangolin species (*Manis javanica*) from which this sequence was obtained could be different from the pangolin species that putatively transferred SARS-CoV-2 to humans (Xiao et al., 2020). A pangolin species with a glycine at position 354 would likely be more vulnerable to a virus similar to SARS-CoV-2.

What can we anticipate about the ACE2 of the true bat reservoir of SARS-CoV-2? We know that MERS-CoV and SARS-CoV-1 receptor orthologs from their respective host species support infection of these viruses with efficiencies similar to or better than their human counterparts (Lau et al., 2018; Li et al., 2005c; Ren et al., 2008; Widagdo et al., 2019). Thus, the ACE2 ortholog of the authentic bat reservoir would be expected to bind the SARS-CoV-2 S protein efficiently. Our ability to identify four residues from ACE2 orthologs of related bats that increase ACE2-Fc binding to this S protein is also consistent with this presumption. If so, we would expect that the reservoir ACE2 would include a glutamic acid at position 24 and a serine at position 34 because both residues enhance ACE2 binding of the SAR2-RBD and they are present in pangolin ACE2 as well. These criteria alone exclude all horseshoe bats with reported ACE2 orthologs except *R. macrotis*. However, *R. macrotis* ACE2 does not efficiently bind the SARS-CoV-2 RBD. Thus either SARS-CoV-2 evolved significantly in a reservoir intermediate, or *R. macrotis* is not itself the most proximal bat reservoir. *R. macrotis* ACE2 includes two unusual residues in the SARS2-RBD-binding region, namely K35 and E42, which are glutamic acid and glutamine, respectively, in most horseshoe-bat orthologs as well as pangolin and human ACE2. We therefore expect that a *bona-fide* SARS-CoV-2 reservoir would include E24, S34, E49, and either E35 or Q42 or both. It is also possible that this true reservoir would, like *R. pusillus*, lack a glycosylation motif at residue 90.

Finally, our studies also identify ways to enhance the neutralization potency of ACE2-Fc. Specifically we show that, although the SARS2-RBD binds human ACE2 with high affinity, it is still imperfectly adapted to this receptor, and its structure still reflects its previous adaptation to its animal hosts. The four potency-enhancing mutations identified here could also be combined with D30E and the ACE2 collectrin domain, both also shown to enhance neutralization by ACE-Fc. Collectively these changes could approach or surpass the potency of most reported SARS-CoV-2 neutralizing antibodies, and thus improve ACE2-Fc, currently under clinical evaluation, as a potential SARS-CoV-2 treatment.

## MATERIALS AND METHODS

### Plasmids

The plasmids for expression of variant coronavirus spike proteins or ACE2 proteins were created by synthesizing fragments by Integrated DNA Technologies (IDT, Coralville, IA, USA), and ligating them into their respective vectors using In-Fusion® HD Cloning Kit (Takara Bio USA) according to manufacturer’s instructions. ACE2-Fc variants were generated by the QuikChange II site-directed mutagenesis protocol (Agilent).

### Cells and viruses

HEK293T (human embryonic kidney; ATCC CRL-3216, VA, USA) were maintained in growth media composed of Dulbecco’s Modified Eagle Medium (DMEM, Life Technologies) supplemented with 2 mM Glutamine (Life Technologies), 1% non-essential amino acids (Life Technologies), 100 U/mL penicillin and 100 µg/mL streptomycin (Life Technologies), and 10% FBS (Sigma-Aldrich, St. Louis, MI, USA) at 37° C in 5% CO2. Expi293 FTM cell (Thermo Fisher) was cultured in Expi293 FTM expression medium (Thermo Fisher) at 37°C in a shaking incubator at 125 rpm and 8% CO_2_.

HEK293T cell line expressing human ACE2 (hACE2) were created by transduction with vesicular stomatitis virus (VSV) G protein-pseudotyped murine leukemia viruses (MLV) containing pQCXIP-myc-hACE2-c9 as previously described (Moore et al., 2004). Briefly, HEK293T cells were co-transfected with three plasmids, pMLV-gag-pol, pCAGGS-VSV-G and pQCXIP-myc-hACE2-c9, and the medium was refreshed after overnight incubation of transfection mix. The supernatant with produced virus was harvested 72h post transfection and clarified by passing through 0.45 μm filter. Parental 293T cells were transduced with generated MLV virus, and the 293T-hACE2 cell lines were selected and maintained with medium containing puromycin (Sigma). hACE2 expression was confirmed by immunofluorescence staining using mouse monoclonal antibody against c-Myc antibody 9E10 (Thermo Fisher) and goat-anti-mouse FITC (Jackson ImmunoResearch Laboratories, Inc). 293T-human ACE2 stable cells were maintained in growth media including 3 μg/ml puromycin for selection of stably transduced cells.

### Protein Production and purification

Expi293 cells (Thermo-Fisher) were transiently transfected using FectoPRO (Polyplus) with plasmids encoding coronavirus RBDs or soluble ACE2 variants with a human-Fc fusion or a C-terminal C-tag (EPEA). After 5 days in shaker culture, media were collected and cleared of debris for 10 min at 1,500×g and filtered using 0.45-µm flasks (Nalgene). Proteins were isolated using MabSelect Sure (GE Lifesciences) or CaptureSelect C-tagXL (Thermo-Fisher) columns according to the manufacturers’ instructions. Eluates were buffer exchanged with PBS and concentrated using Amicon ultra filtration devices (Millipore Sigma) and stored at 4°C before use.

### Flow cytometry to test the binding of coronavirus RBD-Fc proteins to receptors

HEK293T cells were transfected with plasmids encoding ACE2 orthologs or human DPP4 using PEI 40K (Polysciences) according to manufacturer’s instructions. 48h post transfection, transfected cells were detached and incubated with 5 μg/ml RBD-Fc, and the interaction was detected with goat-anti-human-Ig-APC (Jackson ImmunoResearch Laboratories, Inc). Expression of ACE2 orthologs were detected using mouse monoclonal antibody against c-Myc antibody 9E10 (Thermo Fisher) and Goat-anti-mouse FITC (Jackson ImmunoResearch Laboratories, Inc). Samples were analyzed by flow cytometry (BD Accuri C6 Flow Cytometry) and data was analyzed using FlowJo (FlowJo, LLC).

### Surface plasmon resonance studies

Kinetic and thermodynamic parameters for ACE2 binding to RBD-Fc or RBD binding to ACE2-Fc were measured on a Biacore X100 instrument as previously described (Peng et al., 2017). Briefly, a CM5 sensor chip was immobilized with a mouse anti-human IgG CH2 mAb using reagents and instructions supplied with the Human Antibody Capture Kit (GE Healthcare) in order to capture RBD-Fc or ACE2-Fc. Monomeric ACE2 or monomeric RBD serially two-fold diluted in 1x HBS-EP+ running buffer were then injected respectively at five different concentrations with a replicate of the lowest concentration to confirm regeneration of the sensor chip. Calculation of association (*k*_*on*_) and dissociation (*k*_*off*_) rate constants was based on a 1:1 Langmuir binding model. The equilibrium dissociation constant (*K*_*d*_) was calculated from *k*_*off*_*/k*_*on*_.

### Neutralization studies of SARS1-S, SARS2-S or VSV-G pseudotyped MLV viruses

To investigate the inhibition of RBD-Fc proteins, 293T-hACE2 cells were pre-incubated with serially diluted proteins starting from 50 μg/ml. After one-hour incubation at 37°C, pre-treated cells were inoculated with SARS1-S, SARS2-S or VSV-G pseudotyped MLV viruses at MOI=1, and spun at 3000xg, 4°C for 30 min. To compare the neutralization activity of ACE2 proteins, serially diluted proteins were pre-incubated with pseudotype viruses (MOI=1) at 37°C for one hour, and the mixes were added to 293T-hACE2 and spun at 3000×g, 4°C for 30 min. Media was refreshed 2h after further incubation at 37°C after the spin and firefly luciferase activity was measured (Britelight) 48 hours post-infection.

### Computational analysis

IC_50_ analysis was performed after concentration was log_10_ transformed, using default settings for log(inhibitor) vs. response Variable slope method in GraphPad Prism version 8.0.0 for Windows, GraphPad Software, San Diego, California USA, www.graphpad.com.

## ACKNOWLEDGEMENTS

This work is funded by a COVID-19 supplement to AI129868 awarded to M.F. and H.C.

## Author Contributions

H.M., B.D.Q, H.P., C.R., H.C., and M.F. designed the study and its experiments. H.M., B.D.Q, H.P., Y.G., S.P., L.Z., and Z-X.V. performed experiments. M.E.D-G, M.R.G, C.C.B., and M.D.A. provided critical insights and advice, G.C. and H.M. performed statistical analyses. H.M. and M.F. wrote the manuscript with key assistance from B.D.Q., H.P., C.R., and H.C.

## SUPPLEMENTARY MATERIAL FOR

### SUPPLEMENTARY FIGURE LEGENDS

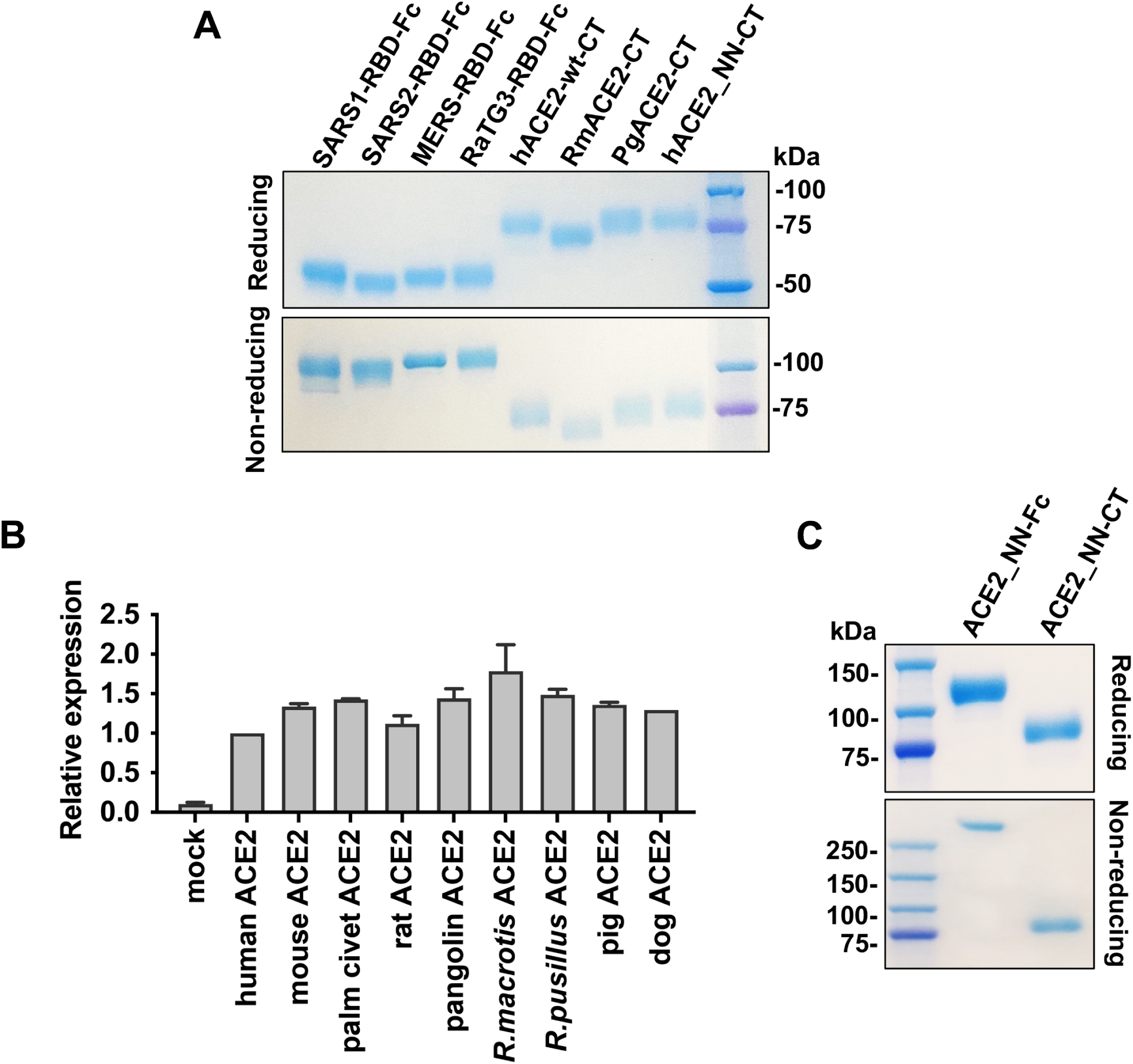
Expression of soluble form RBD-Fc or ACE2 proteins and relative expression of ACE2 orthologs for Figure 1. (**A**) Purified RBD-Fc or monomeric ACE2 proteins were analyzed on NoVEX^®^ 4-20% Tris-Glycine gradient gel under reducing (upper) and non-reducing (lower) conditions. (**B**) HEK293T cells were transfected with control plasmid (pcDNA) or with plasmids expressing ACE2 orthologs and stained by anti-myc 9E10 antibody. The staining was analyzed my flow cytometry. Data are representative of at least three experiments and were performed with those shown in Fig. 1. (**C**) Purified human ACE2-Fc and monomeric human ACE2 proteins were analyzed as in (A). In panels (A) and (C), one microgram of each purified protein was loaded, and the gels were stained with GelCodeBlue reagent. Positions and sizes of the markers are indicated.

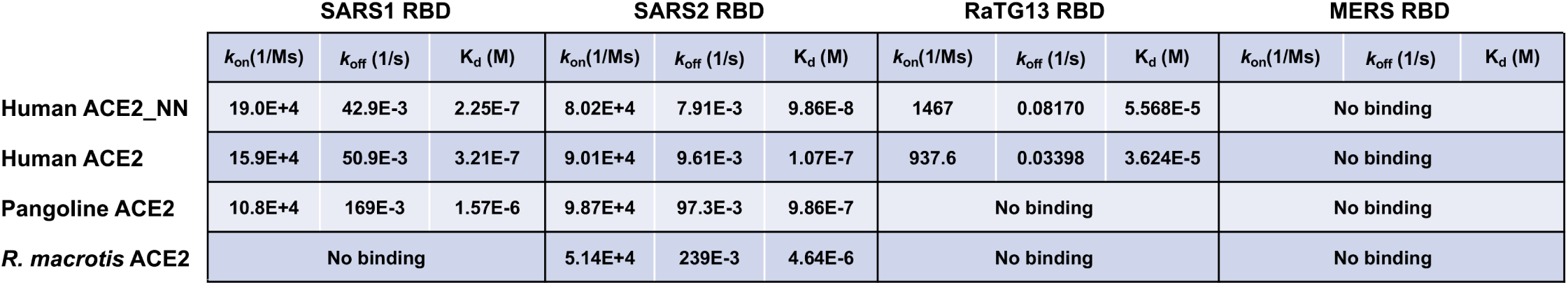
Rate constant values for Figure 2. Association rate constant (*k*_*on*_), dissociation rate constant (*k*_*off*_), and equilibrium dissociation constant (*K*_*d*_) values from SPR analysis of Figure 2.

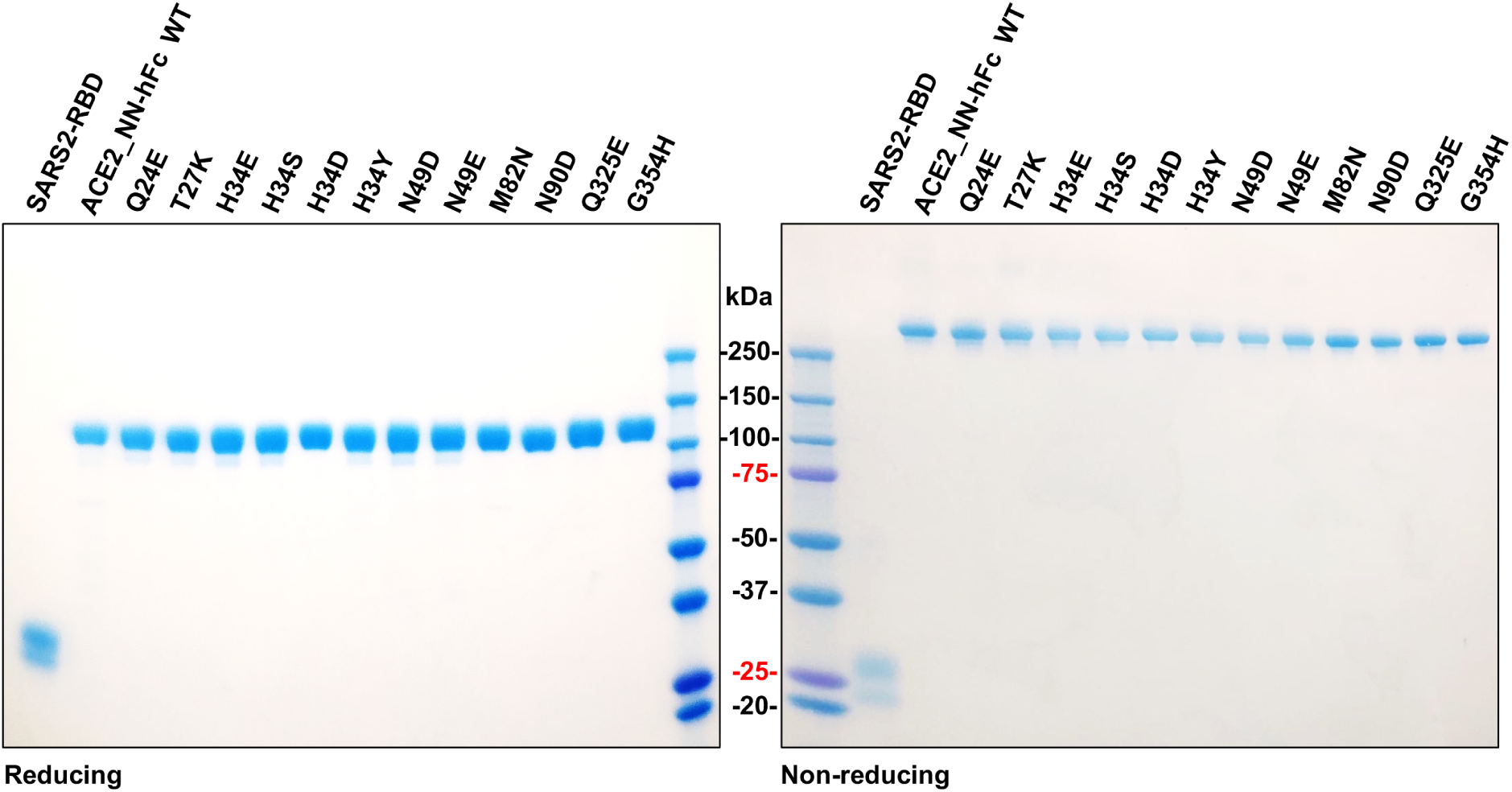
Expression of monomeric SARS2-RBD or ACE2-Fc variants used in Figure 5. Purified monomeric SARS2-RBD or ACE2-Fc variants including any of the single mutation indicated were analyzed under reducing (left) and non-reducing (right) conditions as described in Figure S1.

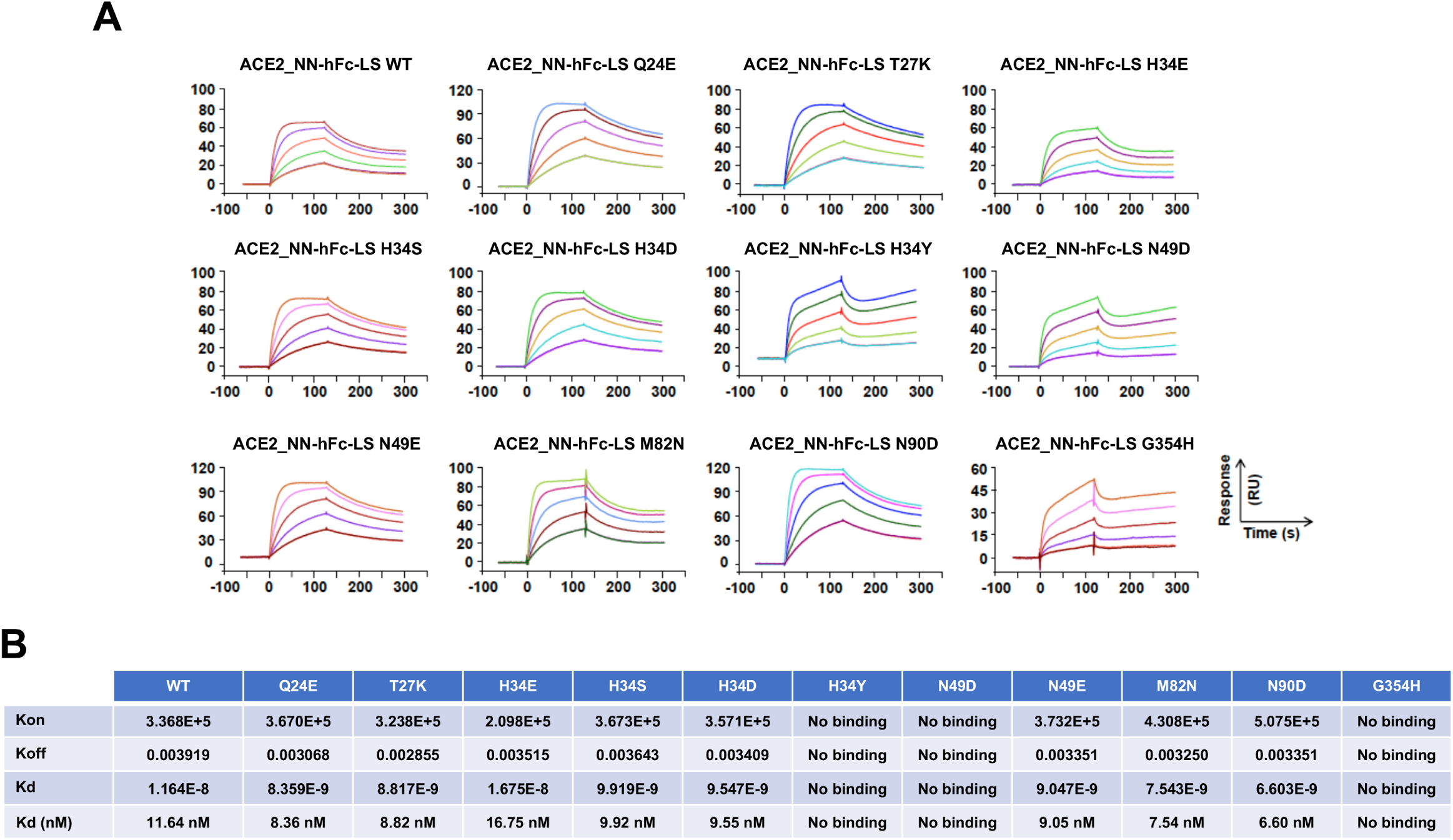
SPR analyses for the binding of monomeric SARS2-RBD to human ACE2-Fc variants, as summarized in Figure 5D. Analysis similar to Figure 2 except that ACE2-Fc variants were captured on the chip and monomeric SARS2-RBD was injected in. (A) Biacore X100 sensorgrams obtained. (B) *k*_*on*_, *k*_*off*_ and *K*_*d*_ from the analysis.

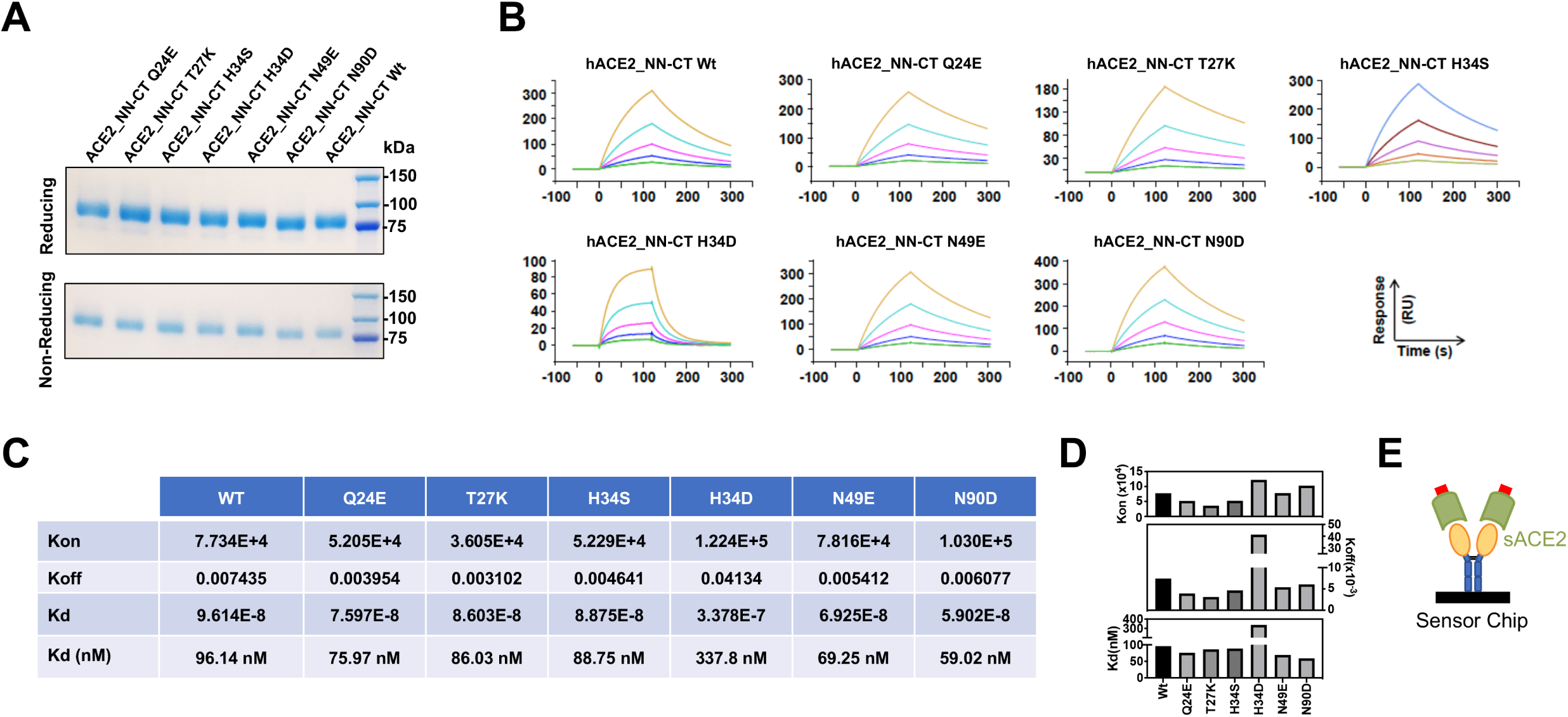
Comparative SPR analyses for the binding of monomeric human ACE2 variants to SARS2-RBD-Fc captured on the chip, as summarized in Figure 6D. (**A**) Purified monomeric ACE2 variants were analyzed on NoVEX^®^ 4-20% Tris-Glycine gradient gel under reducing (upper) and non-reducing (lower) conditions, as in Figure S1. (**B**) The Biacore X100 sensorgrams of the indicated monomeric ACE2 various bound to immobilized SARS2-RBD-Fc are shown. *K*_*on*_, *K*_*off*_ and *K*_*d*_ from this analysis that are presented in a table (**C**) and plotted (**D**). (**E**) A representation of the experiment in (B) indicating that the RBD-Fc was captured and that soluble monomeric ACE2 variants were captured.

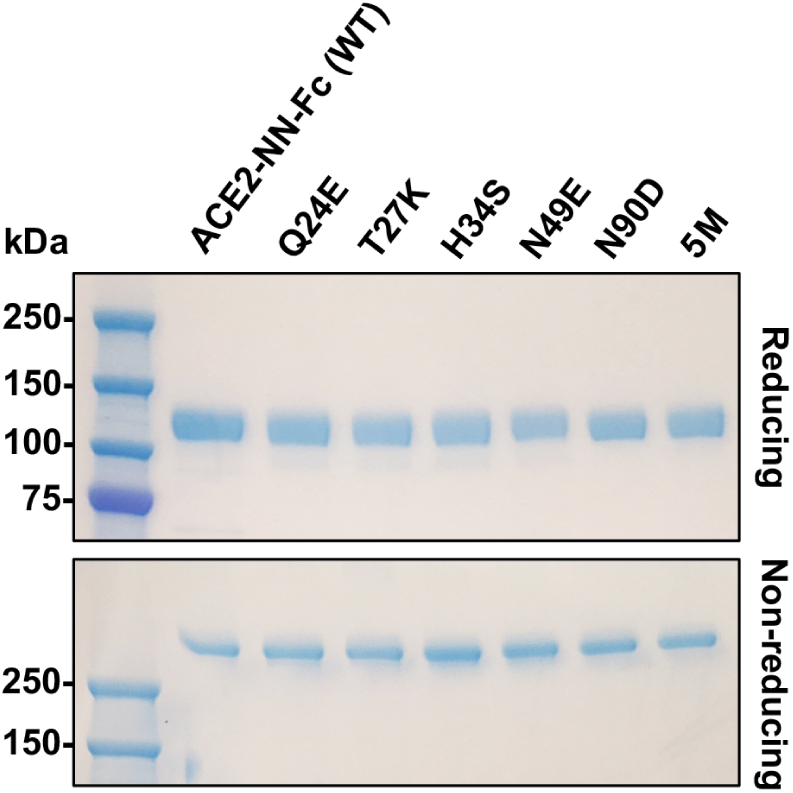
Expression of ACE2-Fc variants used in Figure 6. One microgram of ACE2-Fc variants including either single mutation or all five mutations were analyzed on NoVEX® 4-20% Tris-Glycine gradient gel under reducing (left) and non-reducing (right) conditions.

